# Reporting quality, effect sizes, and biases for aging interventions: a methodological appraisal of the DrugAge database

**DOI:** 10.1101/2025.06.30.660585

**Authors:** Austin Parish, John P.A. Ioannidis, Kevin Zhang, Diogo Barardo, William Swindell, João Pedro de Magalhães

## Abstract

Though interest has grown significantly over the past decades in interventions that may slow the aging process, most evidence for these interventions still comes from experiments in non-human animals. These studies may suffer from design, quality and reporting issues. The quality and reporting of preclinical studies have not yet been studied systematically in anti-aging research. Here we analyzed the DrugAge database, assessing reporting study quality, bias and effect sizes across 667 anti-aging preclinical studies. We found significant shortcomings in reporting of crucial design features such as randomization and blinding, as well as large variation in reporting quality and effects across species. Non-mammal findings typically did not translate to mammals. Although anti-aging interventions may have different effects depending on when they are started, most studies began giving the intervention under investigation very early in the organism’s lifespan. Our findings suggest there is substantial room for improvement in preclinical anti-aging research.

## Introduction

There is increasing interest in interventions targeting the aging process^1,2^. The “geroscience hypothesis” posits that a shared pathophysiology of aging shapes most chronic diseases and interventions targeting aging will confer larger health benefits than those targeting any individual disease^3,4^. Research into such anti-aging interventions has grown substantially, including trials repurposing commonly used drugs such as metformin^5^.

Because of the large sample sizes and long durations of trials required to demonstrate anti-aging effects, most evidence to date has come from preclinical experiments in non-human animals^6^. Aging is a universal pathological process in eukaryotes^7,8^ with conservation of aging pathways across organisms^9,10^; interventions targeting aging may be more successfully translated than interventions for specific diseases which often rely on artificial disease models^11,12^.

Given the possible substantial health benefits of slowing aging, the quality of preclinical studies in this area may be especially important. However, alongside the challenges translating results from one species to another, model organism studies have a long history of shortcomings and design flaws^13,14,15^.

Here, we systematically analyzed studies from DrugAge, a curated database of preclinical experiments investigating the effects of interventions on aging and lifespan in non-human animals^16^. We aimed to evaluate the quality of reporting and methodological rigor of this literature, assess the distribution of observed effect sizes, and probe for the presence of diverse biases. We also investigated how these features changed over time.

## Methods

### Study selection and feature extraction

We downloaded the fifth build of the DrugAge database on May 1^st^, 2025, which contained 3423 different lifespan experiments from a total of 680 unique studies. From these, we excluded 12 studies focusing on replicative aging in the yeast *Saccharomyces cerevisiae* and one duplicate study, yielding 667 unique studies.

For the 32 studies containing experiments with more than one organism, a single experiment was randomly selected for each organism. If a study contained experiments where the same compound was started at different points in the same organism’s lifespan, we included one experiment for the earliest and one experiment for the latest start time. After selecting experiments in this way, our final dataset contained 720 experiments. See **Fig. 1** for a flowchart of data extraction and **Data Supplement 1** for all included studies and experiments. Overall, the 720 experiments represented 568 different species-drug pairs.

**Fig. 1.**
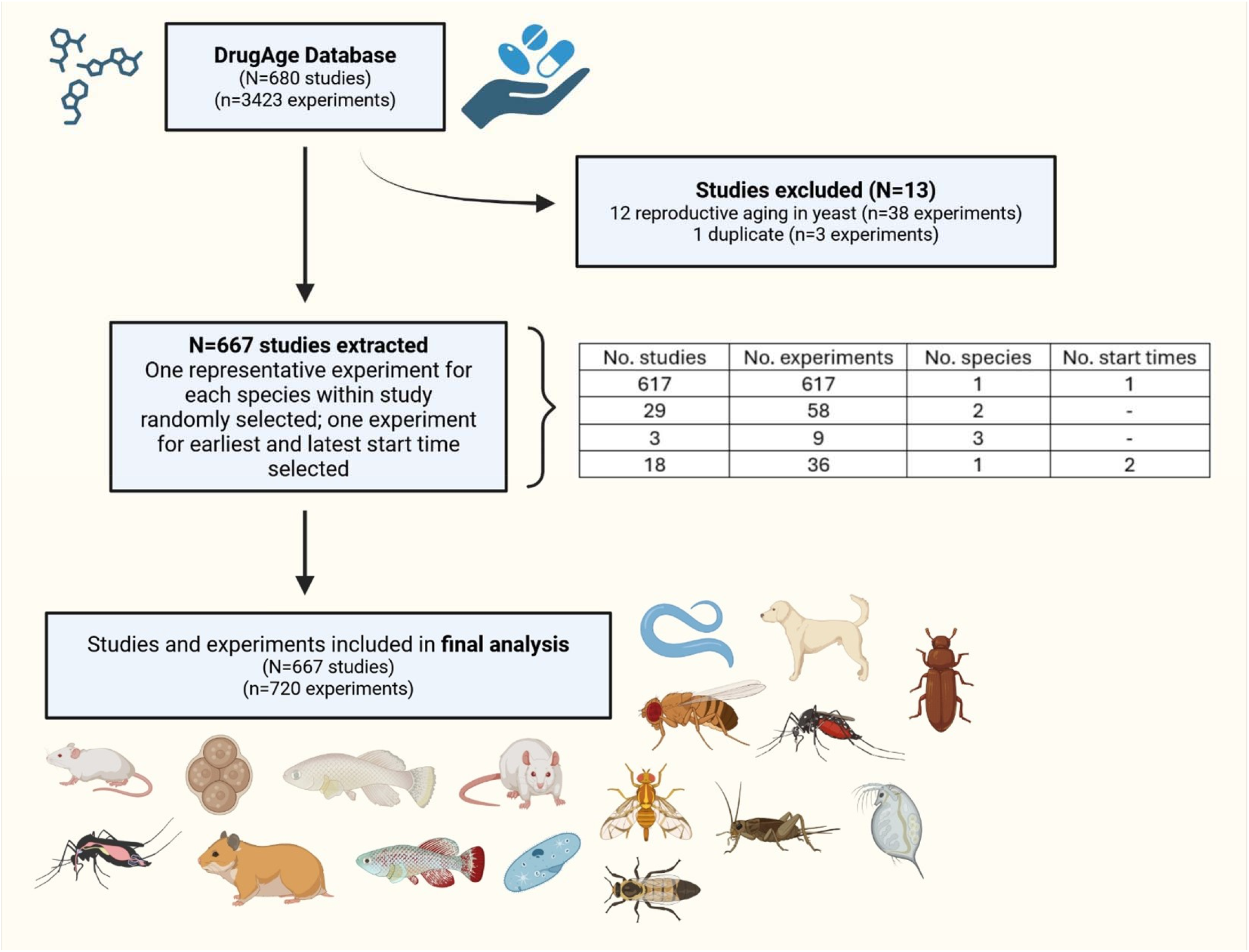
Flowchart of data extraction.

For each study, we extracted eight relevant quality checklist items from CAMARADES (Collaborative Approach to Meta-Analysis and Review of Animal Data in Experimental Studies) (Macleod 2004): 1) whether the study was peer reviewed, and reporting of 2) control of temperature, 3) random allocation to treatment/control; 4) blinded intervention; 5) blinded assessment of outcomes; 6) sample size calculations; 7) adherence to animal welfare regulations; and 8) potential conflicts of interest.

We also extracted the median or mean lifespans of experimental and control groups and whenever possible their confidence intervals (CIs) or standard errors (SEs). When these were not reported, we estimated them from included Kaplan-Meier figures and corresponding log-rank test p-values.

### Statistical techniques

The mean difference between experimental and control groups in lifespan was calculated, as well as the standardized mean difference (SMD) and its SE. Whenever possible, the SEs reported for the experimental and control lifespans were used to calculate the SMD and its SE; when these were not reported, the log-rank p-value was used to estimate the SE^17^. We also calculated the relative increase in lifespan, obtained by dividing the mean difference in lifespan by the average lifespan for that species.

Random-effects meta-analysis was performed using the Sidik-Jonkman estimator^18^. Heterogeneity was estimated with the I^2^ statistic and Q test^19,20^. Meta-analysis calculations used the meta package in R^21^. Contour-enhanced funnel plots and Egger’s test were used to detect small study effects^22^, considering all 720 results together in the same funnel plot. We also applied the test of excess significance and proportion of statistical significance test, tests that may suggest selective reporting biases^23,24^.

Results were considered statistically significant for p-values <0.005 and possibly suggesting significance for p-values between 0.05 and 0.005^25^. The 4.1.0 version of the R programming language was used for all calculations^26^.

## Results

### Quality and design features across studies and species

Of 667 included studies, 617 included only experiments with one species and one start time; 29 summarized experiments with two species and three summarized experiments with three species. Eighteen studies had experiments with two start times, for a total of 720 experiments. Of these, 364 involved an organism that reproduces sexually; of these, 130 used only males (35.7%); 47 used only females (12.9%), 172 used both (47.3%) and 15 did not report the sex(es) used (4.1%). The median sample size across experiments was 200 animals (IQR: 105-338).

All studies were published in peer-reviewed journals (667, 100.0%) and most stated control of temperature (607, 91.0%). Randomization was mentioned in 133 studies (19.9%). Blinding to intervention was mentioned in 27 studies (4.0%), blinded assessment of outcomes in 20 (3.0%), and sample size calculations in 40 (6.0%). Following animal welfare regulations was mentioned in 93 studies (13.9%). Conflict of interest statements were included in 347 studies (52.0%).

The median CAMARADES score across studies was 3 (IQR: 2-3), and varied significantly across species (p<0.0001). 61 studies (9.1%) had CAMARADES counts >4. Except for peer-review publication that was ubiquitous and blinding that was rare, all CAMARADES components varied significantly across species (p<0.0001, **Table 1**). *Caenorhabditis and Drosophila* studies almost always stated control of temperature, but rarely reported randomization or sample size calculations. Studies of mice and rats stated control of temperature less commonly but did better on all other fronts.

**Table 1.**
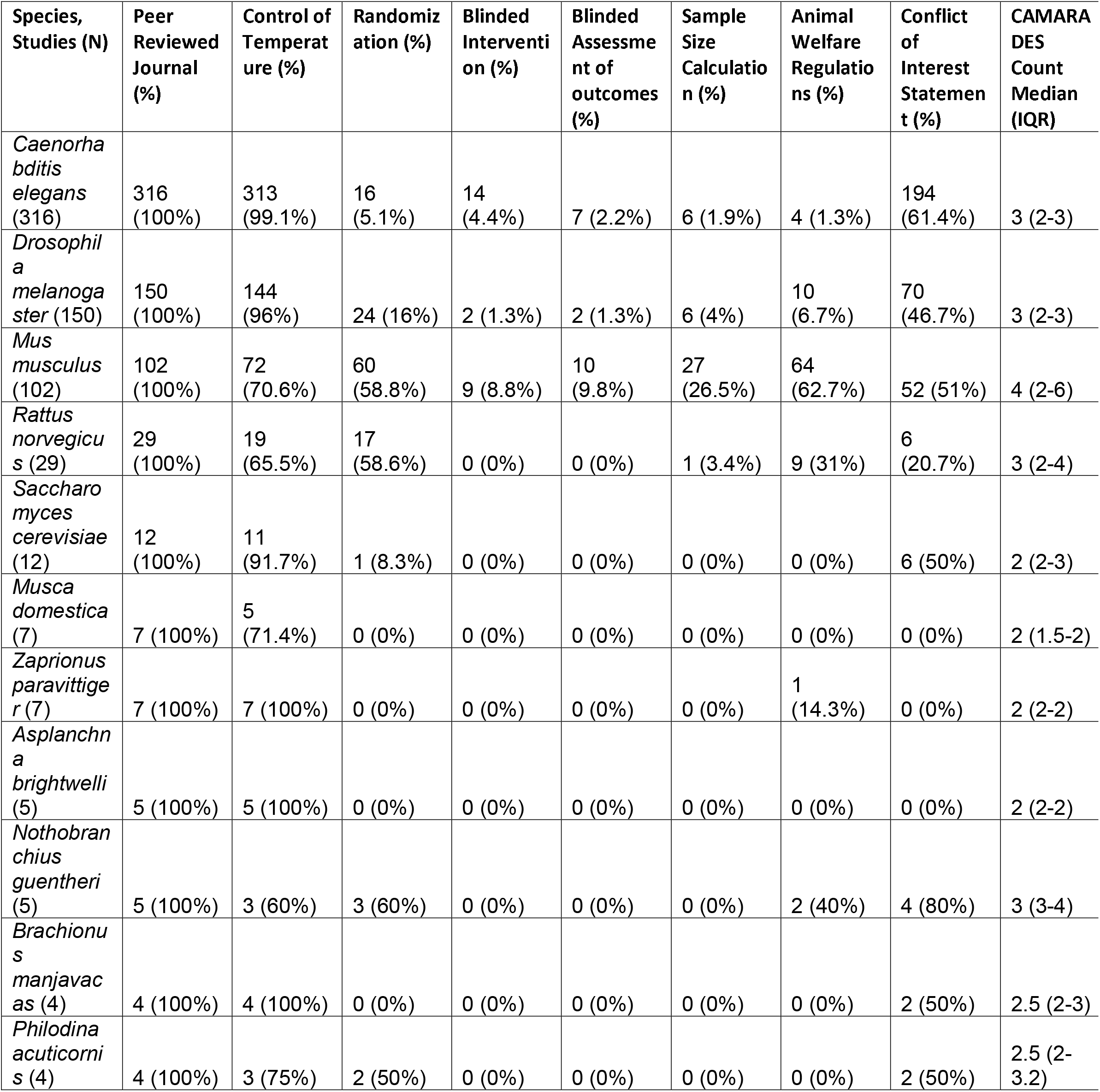
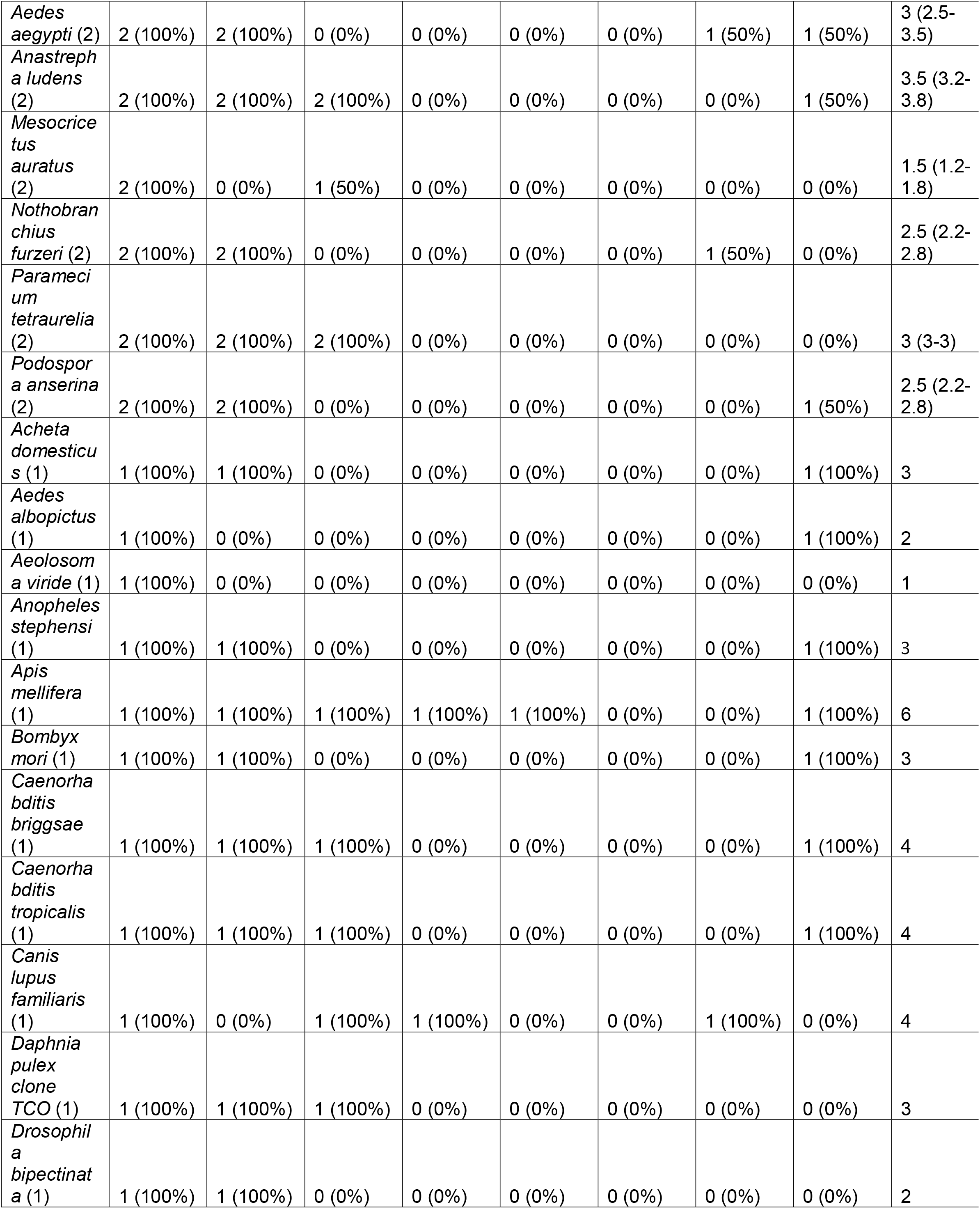
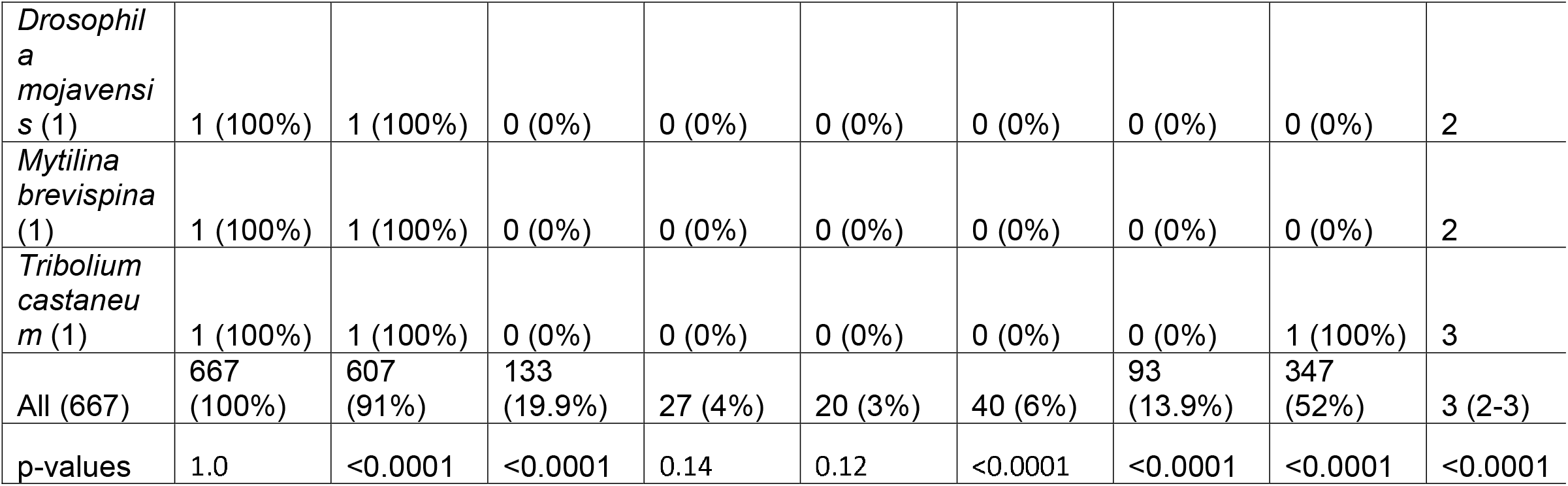
Mention of CAMARADES components across studies of different species in the database, along with median CAMARADES count (sum of the 8 included CAMARADES elements, minimum 0 and maximum 8). P-values for count outcomes reflect the results of 2×31 exact tests; p-value for the median CAMARADES count is the result of the Kruskal-Wallis test.

Of the 667 studies, 153 reported whether the organisms included were from an inbred (genetically homogenous) line or an outbred/hybrid line; of these, 73 used inbred lines. None of the CAMARADES components differed significantly between inbred and non-inbred studies.

### Change in Features Over Time

The earliest included study was published in 1948, the latest in 2024. Over time, there was a significant increase in reporting conflicts of interest, compliance with animal welfare regulations, control of temperature and sample size calculations (linear regression p<0.0001 for each). There was no significant increase over time in reporting of randomization (p=0.60), or blinding with regard to intervention (p=0.07) or outcomes (p=0.011) (**Fig. 2**). Studies had higher CAMARADES counts over time (p<0.0001).

**Fig. 2.**
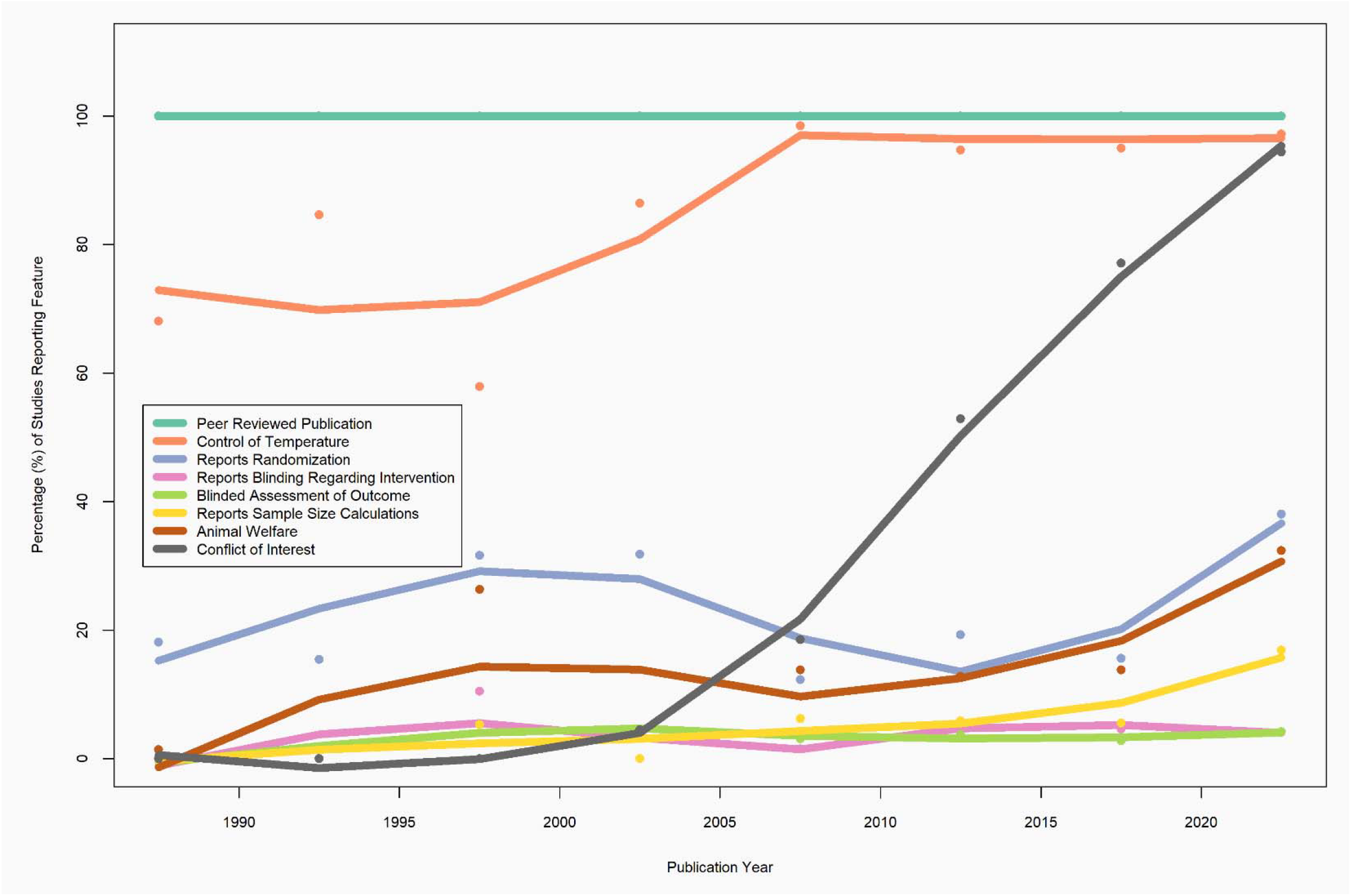
Percentage of studies with specific CAMARADES features over time (scatterplot, with local polynomial regression fit curves). Reporting of potential conflicts of interest, animal welfare regulations, control of temperature and sample size calculations increased significantly over time (p<0.0001 for each).

### Average Start Time as Percentage of Organism Average Lifespan

Across the 720 experiments, the median percentage of average lifespan that interventions were started at was 6.0% (IQR: 4.3-12.8%) (**Fig. 3**), with significant variation across species (**Table 2**). Mammal experiments started at a relatively later point in lifespan than non-mammal experiments (25.6% vs 5.9%, p<0.0001). Most experiments started “early” (before 20% of average lifespan) (n = 596, 82.8%). Few experiments started at 50% of average lifespan or later (52, 7.2%).

**Table 2.** Timepoint in lifespan of organisms that each experiment was started at expressed as a percentage of the median lifespan for that organism. Kruskal-Wallis test for comparison between species p<0.0001.

| Species | Number of experiments | Percentage of average lifespan start time (median, IQR) |
| --- | --- | --- |
| *Caenorhabditis elegans* | 333 | 6.0% (6-6) |
| *Drosophila melanogaster* | 159 | 1.4% (1.4-2.9) |
| *Mus musculus* | 117 | 32.9% (11-60.7) |
| *Rattus norvegicus* | 32 | 14.4% (7.4-36.7) |
| *Saccharomyces cerevisiae* | 13 | 10.0% (10-10) |
| *Musca domestica* | 8 | 3.6% (3.6-3.6) |
| *Zaprionus paravittiger* | 7 | 3.8% (3.8-3.8) |
| *Asplanchna brightwellii* | 5 | 20.0% (20-20) |
| *Nothobranchius guentheri* | 5 | 28.7% (28.7-28.7) |
| *Brachionus manjavacas* | 4 | 8.3% (6.6-25) |
| *Philodina acuticornis* | 4 | 2.6% (2.6-2.6) |
| *Caenorhabditis briggsae* | 3 | 4.9% (4.9-4.9) |
| *Mesocricetus auratus* | 3 | 6.6% (4.7-24.6) |
| *Aedes aegypti* | 2 | 38.1% (21.4-54.8) |
| *Anastrepha ludens* | 2 | 1.4% (0.8-1.9) |
| *Caenorhabditis tropicalis* | 2 | 5.6% (5.6-5.6) |
| *Nothobranchius furzeri* | 2 | 11.1% (10.4-11.7) |
| *Paramecium tetraurelia* | 2 | 1.8% (1.8-1.8) |
| *Podospora anserina* | 2 | 10.9% (7.6-14.1) |
| *Acheta domesticus* | 1 | 11.7% |
| *Adineta vaga* | 1 | 12.6% |
| *Aedes albopictus* | 1 | 2.5% |
| *Aeolosoma viride* | 1 | 1.7% |
| *Anopheles stephensi* | 1 | 25% |
| *Apis mellifera* | 1 | 11.1% |
| *Bombyx mori* | 1 | 13.3% |
| *Canis lupus familiaris* | 1 | 74.1% |
| *Daphnia pulex* clone TCO | 1 | 29.0% |
| *Drosophila bipectinata* | 1 | 4.3% |
| *Drosophila kikkawai* | 1 | 4.0% |
| *Drosophila mojavensis* | 1 | 40.7% |
| *Drosophila virilis* | 1 | 1.4% |
| *Mytilina brevispina* | 1 | 34.5% |
| *Tribolium castaneum* | 1 | 1.2% |
| All species | 720 | 6.0% (4.3-12.8) |

**Fig. 3.**
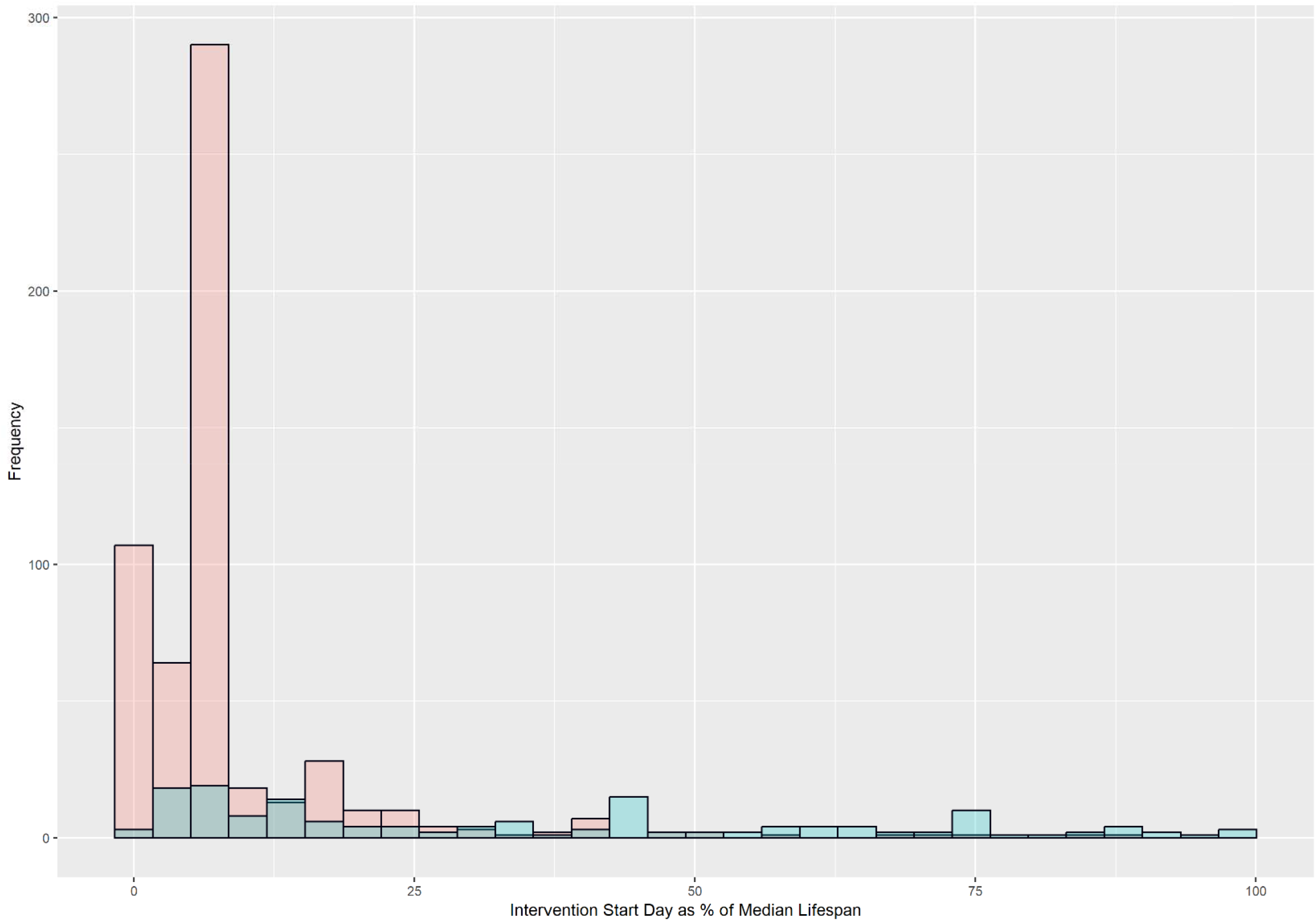
Distribution of when an intervention was started in the lifespan of an organism, expressed as a percentage of the median lifespan for that species, across 720 experiments. Blue represents 153 mammal experiments (median = 25.6%), and pink represents 567 non-mammal experiments (median = 6.0%).

### Distribution of Effect Sizes

Of the 720 included experiments, most SMDs were positive (638, 88.6%), indicating a favorable effect of the intervention on lifespan. The median SMD was 0.43 (IQR: 0.24-0.70); the random effects meta-analysis estimate was 0.57 (95% CI: 0.48-0.66, p<0.0001), with significant heterogeneity (I = 95%, 94-96%, p<0.0001). As a fraction of average species lifespan, the median percentage increase in lifespan was 11.4% (IQR: 5.4-19.1%); the meta-analysis estimate was 12.2% (11.0-13.4%, p<0.0001). **Table 3** summarizes these results.

**Table 3.**
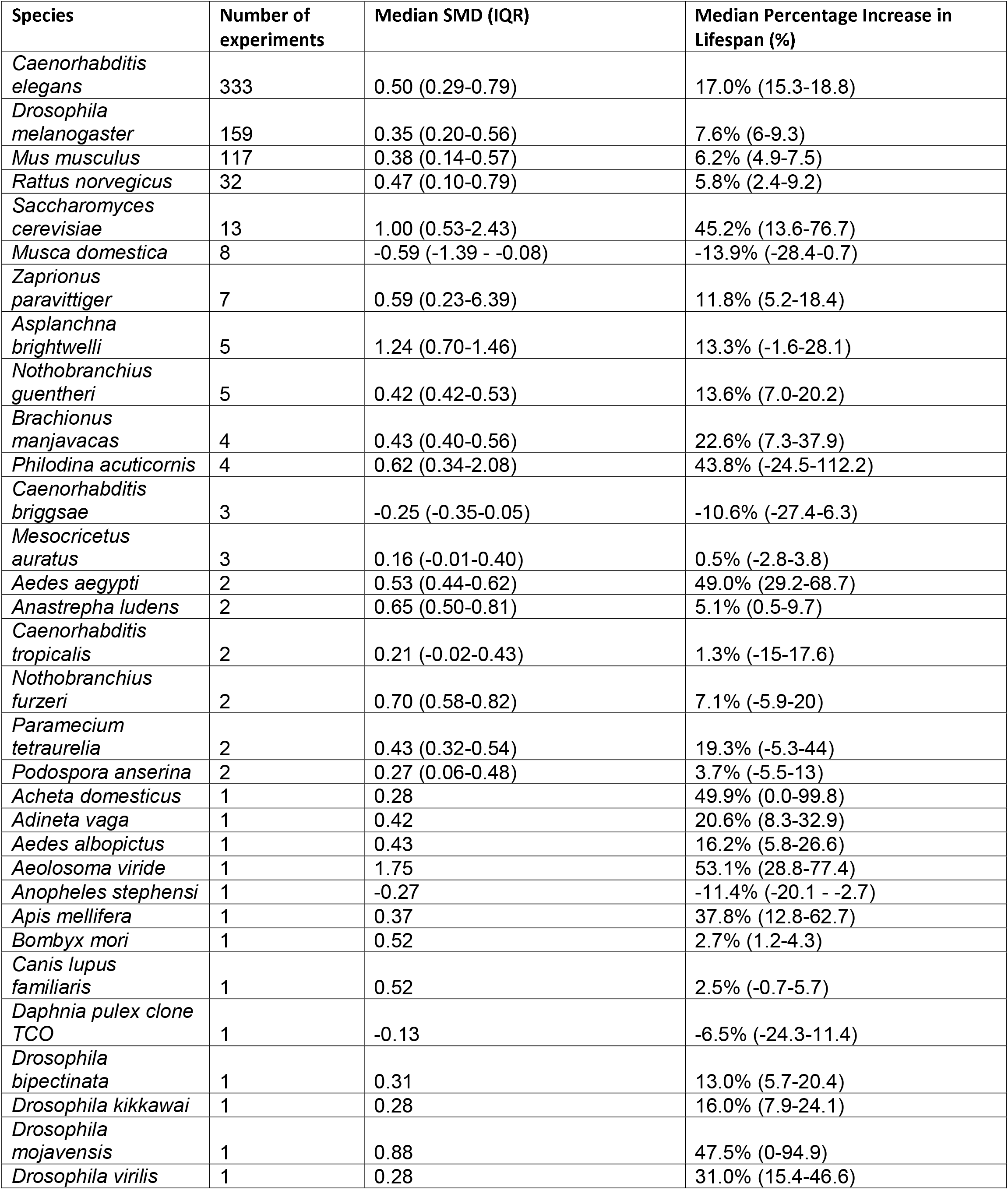
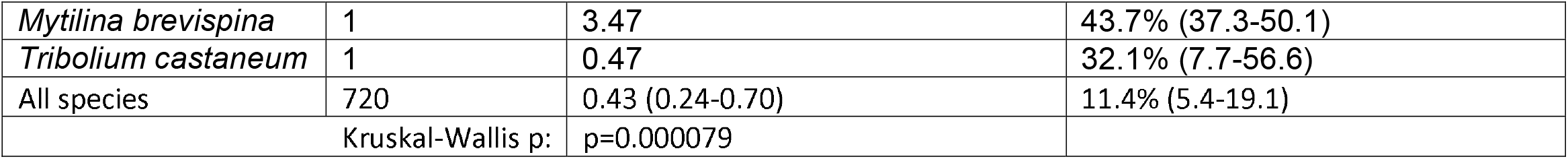
Median and IQR of SMD for lifespan increase, as well as the median percentage increase in lifespan, for each species in the database, across 720 experiments from 667 studies.

| Species | Number of experiments | Median SMD (IQR) | Median Percentage Increase in Lifespan (%) |
| --- | --- | --- | --- |
| *Caenorhabditis elegans* | 333 | 0.50 (0.29-0.79) | 17.0% (15.3-18.8) |
| *Drosophila melanogaster* | 159 | 0.35 (0.20-0.56) | 7.6% (6-9.3) |
| *Mus musculus* | 117 | 0.38 (0.14-0.57) | 6.2% (4.9-7.5) |
| *Rattus norvegicus* | 32 | 0.47 (0.10-0.79) | 5.8% (2.4-9.2) |
| *Saccharomyces cerevisiae* | 13 | 1.00 (0.53-2.43) | 45.2% (13.6-76.7) |
| *Musca domestica* | 8 | -0.59 (-1.39 - -0.08) | -13.9% (-28.4-0.7) |
| *Zaprionus paravittiger* | 7 | 0.59 (0.23-6.39) | 11.8% (5.2-18.4) |
| *Asplanchna brightwelli* | 5 | 1.24 (0.70-1.46) | 13.3% (-1.6-28.1) |
| *Nothobranchius guentheri* | 5 | 0.42 (0.42-0.53) | 13.6% (7.0-20.2) |
| *Brachionus manjavacas* | 4 | 0.43 (0.40-0.56) | 22.6% (7.3-37.9) |
| *Philodina acuticornis* | 4 | 0.62 (0.34-2.08) | 43.8% (-24.5-112.2) |
| *Caenorhabditis briggsae* | 3 | -0.25 (-0.35-0.05) | -10.6% (-27.4-6.3) |
| *Mesocricetus auratus* | 3 | 0.16 (-0.01-0.40) | 0.5% (-2.8-3.8) |
| *Aedes aegypti* | 2 | 0.53 (0.44-0.62) | 49.0% (29.2-68.7) |
| *Anastrepha ludens* | 2 | 0.65 (0.50-0.81) | 5.1% (0.5-9.7) |
| *Caenorhabditis tropicalis* | 2 | 0.21 (-0.02-0.43) | 1.3% (-15-17.6) |
| *Nothobranchius furzeri* | 2 | 0.70 (0.58-0.82) | 7.1% (-5.9-20) |
| *Paramecium tetraurelia* | 2 | 0.43 (0.32-0.54) | 19.3% (-5.3-44) |
| *Podospira anserina* | 2 | 0.27 (0.06-0.48) | 3.7% (-5.5-13) |
| *Acheta domesticus* | 1 | 0.28 | 49.9% (0.0-99.8) |
| *Adineta vaga* | 1 | 0.42 | 20.6% (8.3-32.9) |
| *Aedes albopictus* | 1 | 0.43 | 16.2% (5.8-26.6) |
| *Aeolosoma viride* | 1 | 1.75 | 53.1% (28.8-77.4) |
| *Anopheles stephensi* | 1 | -0.27 | -11.4% (-20.1 - -2.7) |
| *Apis mellifera* | 1 | 0.37 | 37.8% (12.8-62.7) |
| *Bombyx mori* | 1 | 0.52 | 2.7% (1.2-4.3) |
| *Canis lupus familiaris* | 1 | 0.52 | 2.5% (-0.7-5.7) |
| *Daphnia pulex clone TCO* | 1 | -0.13 | -6.5% (-24.3-11.4) |
| *Drosophila bipectinata* | 1 | 0.31 | 13.0% (5.7-20.4) |
| *Drosophila kikkawai* | 1 | 0.28 | 16.0% (7.9-24.1) |
| *Drosophila mojavensis* | 1 | 0.88 | 47.5% (0-94.9) |
| *Drosophila virilis* | 1 | 0.28 | 31.0% (15.4-46.6) |
| *Mytilina brevispina* | 1 | 3.47 | 43.7% (37.3-50.1) |
| *Tribolium castaneum* | 1 | 0.47 | 32.1% (7.7-56.6) |
| All species | 720 | 0.43 (0.24-0.70) | 11.4% (5.4-19.1) |
| Kruskal-Wallis p: |  | p=0.000079 |  |

Comparing experiments in studies with specific CAMARADES components, reporting of randomization was associated with a smaller SMD (0.38 in those reporting vs 0.45, Kruskal-Wallis p=0.0074). Other CAMARADES components were not associated with significant differences: peer-reviewed publication (p=1.0), control of temperature (p=0.094), blinded intervention (p=0.35), blinded assessment of outcome (p=0.84), sample size calculations reported (p=0.17), compliance with animal welfare regulations (p=0.48), conflict of interest statement (p=0.041).

There was no significant difference between the 596 early start experiments and the 124 late start experiments (median SMD 0.43 vs 0.41, Kruskal-Wallis p=0.46). Median SMD did not vary significantly with publication year (p=0.11) (see **Supplementary Fig. 1** for bubble plot). Studies with mammalian species had lower median SMDs than non-mammal studies (0.39 vs 0.44, p=0.040).

**Supplementary Fig. 1.**
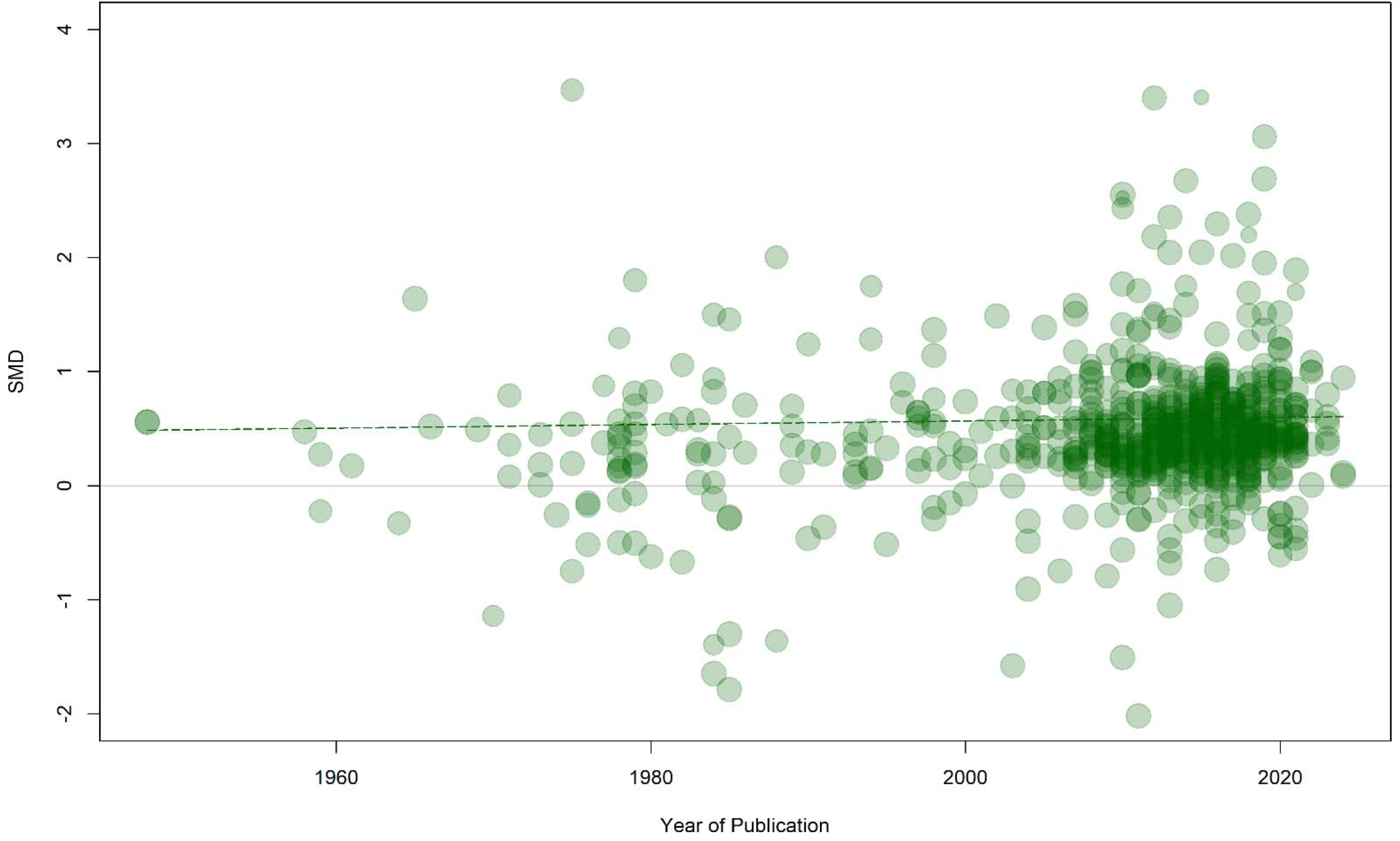
Bubble plot of SMD versus year of publication, for 720 experiments across 667 studies. Diameter of bubbles is proportional to inverse variance of SMD, with larger bubbles representing smaller variance. A linear regression line of best fit is included.

There were 36 compounds that were tested in at least one mammal and at least one non-mammal experiment, allowing for comparisons within the same drug (see **Supplementary Table 1**). Of these, 22 showed a significant increase in lifespan (p<0.005) for non-mammals. Of these, only 8 also showed a significant increase in mammal lifespan (curcumin, spermidine, epithalamin, D-glucosamine, estradiol, simvastatin, SKQ and taurine); additionally, two in contrast showed a significant decrease in mammal lifespan (quercetin and butylated hydroxytoluene).

**Supplementary Table 1.**
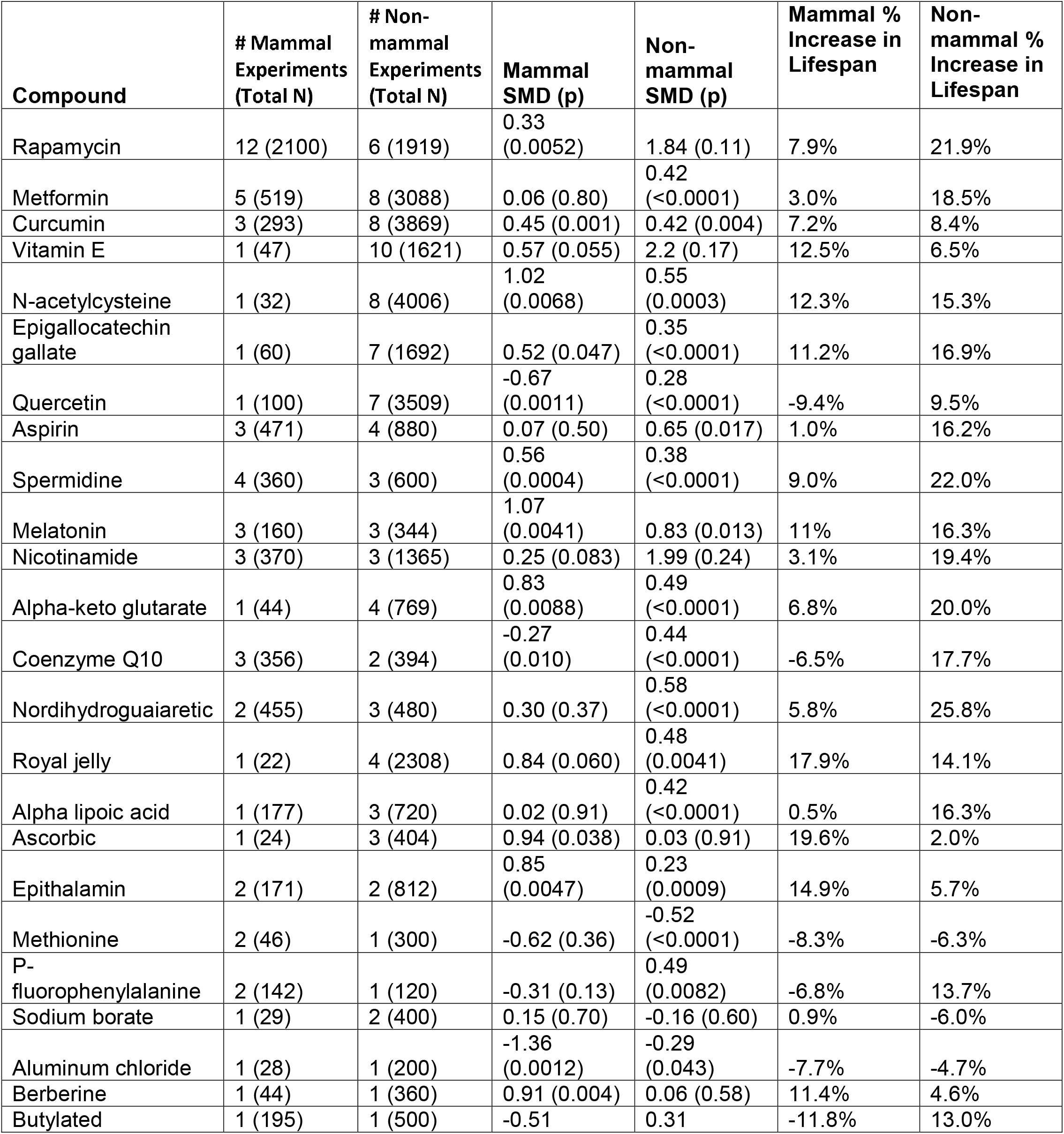
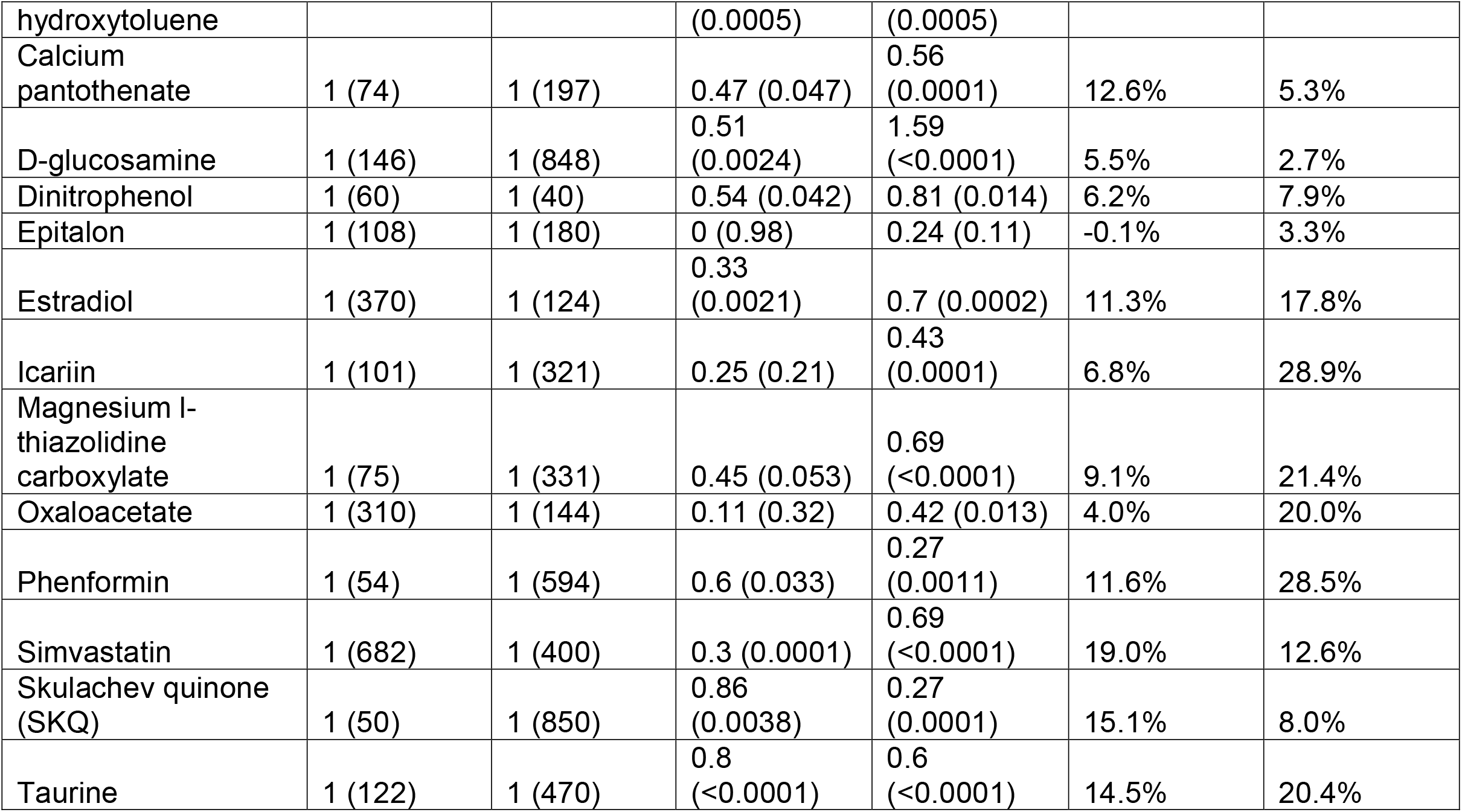
Comparison between mammal and non-mammal experimental results in the 36 compounds with both mammal and non-mammal experiments.

Ten compounds showed a significant increase in mammal lifespan (p<0.005) and most showed some increase in non-mammals as well (p<0.005 for eight of them, as above). However, the amount of mammalian evidence for these compounds was limited (total sample sizes: 293 for curcumin, 360 for spermidine, 160 for melatonin, 171 for epithalamin, 44 for berberine, 146 for D-glucosamine, 370 for estradiol, 682 for simvastatin, 50 for SKQ and 122 for taurine).

The absolute percent error between non-mammal and mammal effects was 78% (IQR: 49-163%). Across compounds, the median percentage increase in mammal lifespan was significantly smaller than the percentage increase in non-mammal lifespan (7.0% vs 14.7%, paired Wilcoxon p=0.004). There was no significant linear correlation between non-mammal and mammal SMDs or percentage increases (r = 0.19, p = 0.30 and r = 0.28, p = 0.11 respectively).

### Distribution of p-values

Of the 720 experiments, 638 (88.6%) were associated with an increase in lifespan and 82 (11.4%) were associated with a decrease in lifespan. Of the 638 experiments associated with increasing lifespan, 495 showed p-values <0.05 (77.6%); of the 82 experiments associated with decreasing lifespan, 51 showed p-values <0.05 (62.2%); **Supplementary Fig. 2** shows the p-value distributions.

**Supplementary Fig. 2.**
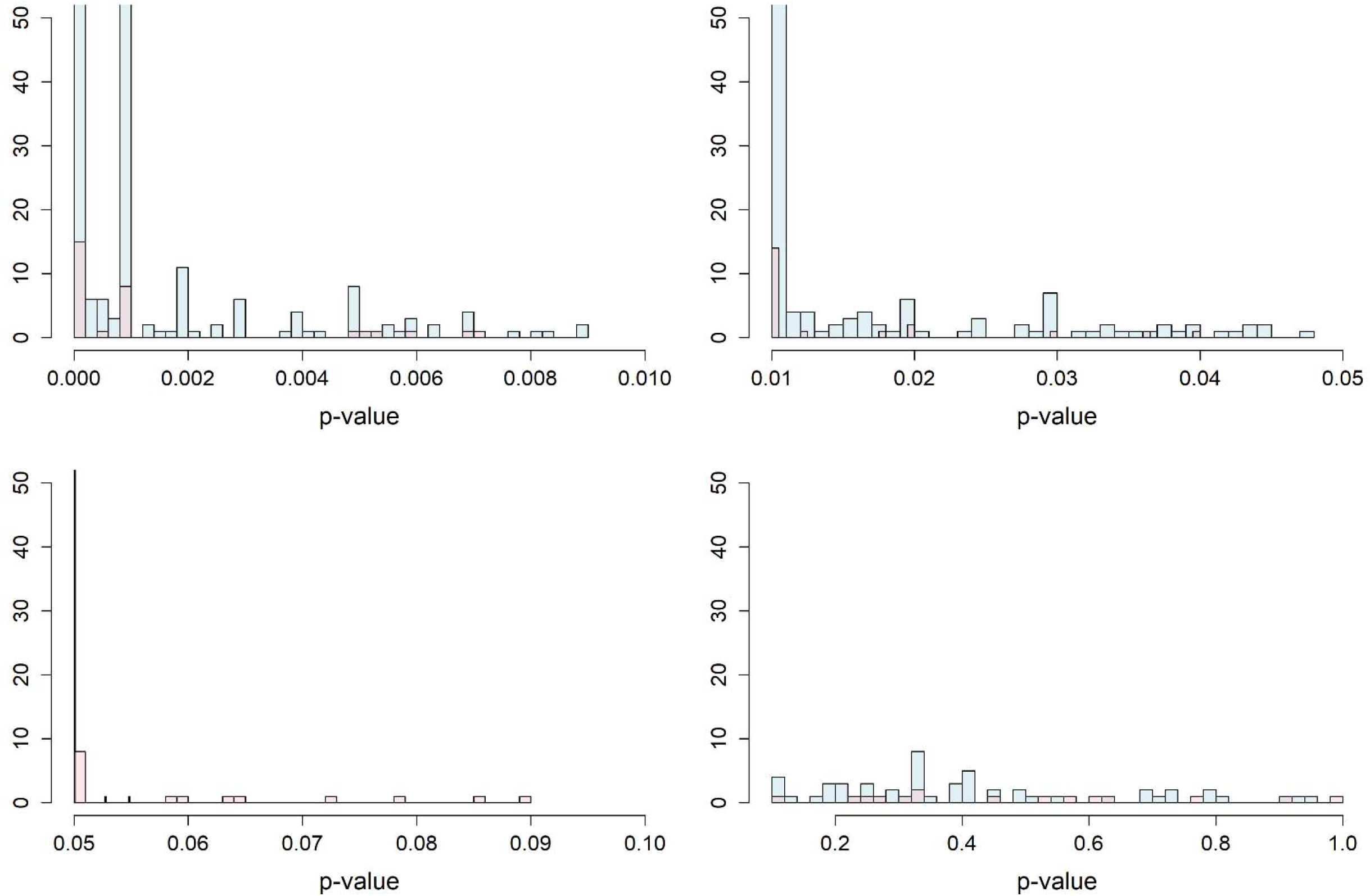
Histogram of p-values from 720 experiments where p-values could be estimated. Blue represents p-values associated with increase in lifespan (positive SMDs) and pink represents p-values associated with decrease in lifespan (negative SMDs). Clockwise from upper left: 0<p<0.01; 0.01<p<0.05; 0.05<p<0.10; 0.10<p<1.0.

### Funnel plot asymmetry and excess significance testing

Across the 720 experiments studies there was evidence of significant funnel plot asymmetry (Egger’s Z = 11.3, p<0.0001); see **Fig. 4** for the contour enhanced funnel plot. The expected number of significant findings was 499 out of 720, while the observed number was 546, indicating significantly more significant results than expected (test of excess significance χ^2^ = 14.0, p<0.0001); the proportion of statistical significance test resulted in a test statistic of Z = 3.74 (p<0.0001)^23,24^.

**Fig. 4.**
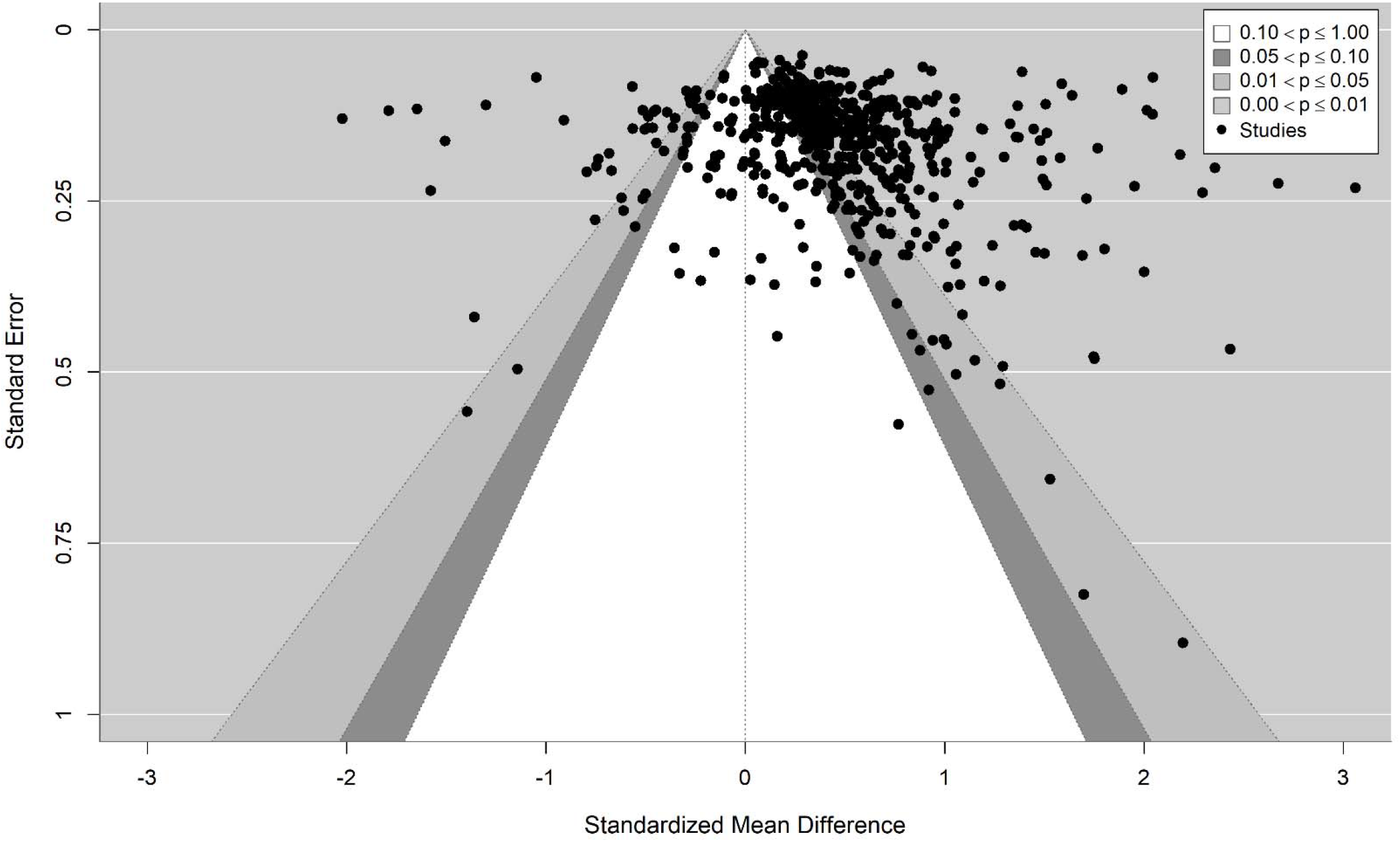
Contour-enhanced funnel plot of SMD from the 720 experiments.

## Discussion

Despite growing excitement about the possibility of anti-aging interventions impacting human healthspan and lifespan, most studies of these interventions have been conducted in non-human animals. In this review of 720 experiments from 667 such studies, we found widely varying reporting of study quality, design and effect sizes across species and compounds. Important design features such as randomization, blinding of intervention, blinded assessment of outcome, compliance with animal welfare regulations, and sample size calculations were infrequently reported, despite evidence that the absence of such features can bias experimental results^27,28,29,30,31^. Only slightly more than half included conflict of interest statements, although all studies were published in peer-reviewed journals and over 90% reported control of temperature.

Although reporting quality improved somewhat over time, this was mainly due to increases in reporting of compliance with animal welfare regulations or conflict of interest statements; crucial design features such as randomization and blinding did not increase substantially over time. Generally, most studies did not meet standard reporting guidelines for pre-clinical experiments^30^.

Preclinical studies on various diseases have also shown infrequent reporting of randomization and blinding. A review of 271 preclinical studies across different diseases found 13% of studies reported randomization and 14% blinding^32^. In another similar review of 290 studies, 32% reported randomization and 11% blinding^33^. Overall, our results are comparable to these, although the reporting of both randomization and blinding seems to be even less frequent in the DrugAge database.

In addition to the overall low rate of reporting, we found significant differences across species: the four most represented species in the database (nematodes, fruit flies, mice and rats) varied widely in reporting of randomization and blinding, as well as in the average effect size found. Over half of mammal studies in the database reported randomization, while less than 10% of non-mammal studies did. The better reporting quality of mammal studies does not alleviate concerns, since even for mammal studies reporting was often suboptimal and most studies in the database were from non-mammals. Additionally, the average effect found in non-mammal studies was significantly larger than that found in mammal studies.

For 36 compounds with both mammal and non-mammal experiments, only eight showed a significant lifespan increase in both non-mammals and mammals; the number of experiments and sample sizes for these results were limited. These results are exploratory, and the numbers are small, but they raise hesitation about the direct translation of these results to more complex organisms such as humans.

Furthermore, previous work has suggested that some interventions may have different effects if started late in an organism’s lifespan rather than early^34,35^, and there is significant interest in discovering interventions that slow aging in older adults^36^. In our assessment, we found that most preclinical experiments started the anti-aging intervention early in the organism’s lifespan, often prior to sexual maturity, when key senescence mechanisms may lack relevance^37^. Although we did not find a significant difference in the effect of interventions between early and late start experiments, the sparsity of late start results makes this comparison uncertain. Our study clearly highlights the paucity of late start experiments in the literature, a deficit of evidence that needs to be remedied. We emphasize the need for whole-lifespan aging experiments with a greater diversity of start times, including more starting in middle or late life, as these better reflect the intended translational application of anti-aging interventions and likely design of future clinical trials investigating proposed interventions.

Overall, the analyzed studies have a plethora of significant reported results and many studies suggest sizeable effect sizes. However, the lack of methodological rigor (at least based on reported information) and the strong prese suggestion of bias (larger effects in smaller studies and an excess of significant results) prompt skepticism about this overall favorable picture and prospects for translation to humans.

### Limitations

Our work has several limitations. The DrugAge database may not include some compounds that have never shown any promising results. Moreover, we did not extract quantitative data from all of the 3423 experiments in the database, but rather extracted a random experiment from each species represented in each study, as well as the earliest and latest start time experiments, obtaining quantitative data from only approximately 21% of the experiments in the database. Although experiments were selected randomly, this method may still have led to a biased estimate of quantitative effects in some cases. Nevertheless, our selection process resulted in a dataset with largely independent observations, while the full database may have a lot of highly correlated data and may have over-represented specific experiments that were rather similar.

It is also possible that some studies that did not mention randomization still carried out randomization, and the same may apply to other design features. Nevertheless, the large variation in reporting of randomization and blinding is concerning. Also, we found the reporting of randomization did not differ significantly between studies that used genetically homogenous populations of organisms and those using more heterogenous populations. Furthermore, while we observed improvements over time in compliance with animal welfare, control of temperature and sample size calculations, it is unclear whether this represents genuine improvements on what studies did, or simply better realization that there are features that should be reported in their publications.

Finally, our analyses were built upon the DrugAge database, which although meticulously constructed may still contain some inaccuracies. Moreover, single studies may have reported some inaccurate results, and we also noted some studies with potentially spurious numbers, e.g., a confusion between standard deviation and standard errors is not rare in studies with continuous outcomes. Nevertheless, data inaccuracies are unlikely to be large enough to invalidate the big picture described by our analyses.

## Conclusion

Preclinical experiments investigating anti-aging interventions do not regularly follow reporting guidelines and infrequently report important design features such as randomization and blinding. There are significant differences in the average lifespan effect, as well as study quality, across different species commonly used in preclinical experiments. Non-mammal results do not seem to reliably predict mammal results, raising further concern for translation. Despite the interest in interventions able to slow aging when initiated late in human lifespan, most preclinical experiments started interventions early in organism lifespans. Our work highlights multiple concrete areas for improvement of preclinical anti-aging research, areas that may be critical for successful translation into human trial results.

## Author Contributions

Austin Parish conceived and designed the study, collected data, performed analysis, and wrote and edited manuscript. John Ioannidis supported the analysis, wrote and edited manuscript.

Kevin Zhang collected data and wrote and edited manuscript.

Diogo Barardo supported the analysis, wrote and edited manuscript. William Swindell supported the analysis, wrote and edited manuscript.

João Pedro de Magalhães supported the analysis, wrote and edited manuscript.

## Acknowledgements

The authors have no acknowledgements to list.

