## Supplementary Tables and Figures for "Reporting quality, effect sizes, and biases for aging interventions: a methodological appraisal of the DrugAge database": DrugAge Aging Quality Review Draft 6-30-25 NA Supplementary Files.pdf

**Supplementary Figures and Tables**

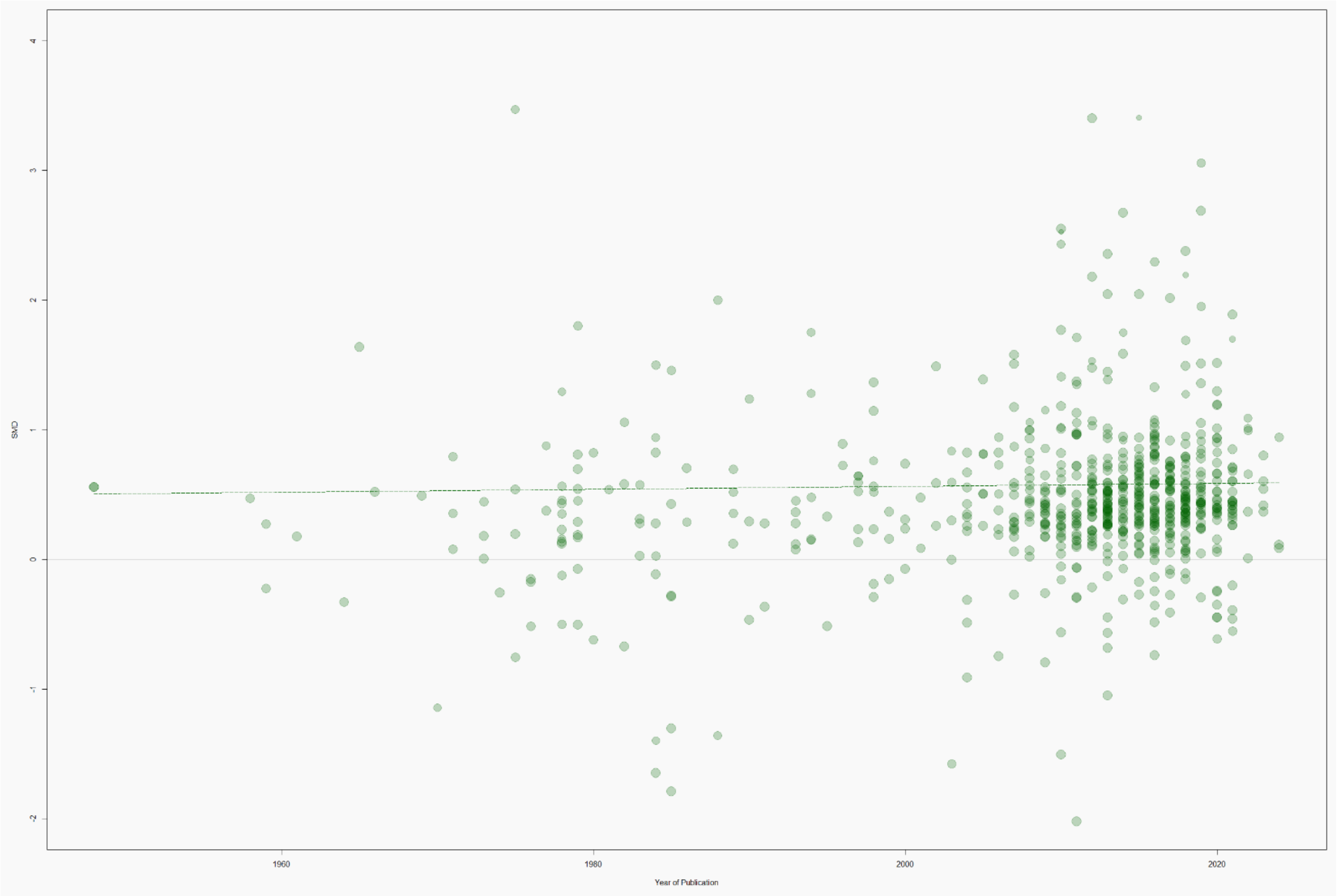


**Supplementary Figure 1.** Bubble plot of SMD versus year of publication, for 720 experiments across 667 studies. Diameter of bubbles is proportional to inverse variance of SMD, with larger bubbles representing smaller variance. A linear regression line of best fit is included.

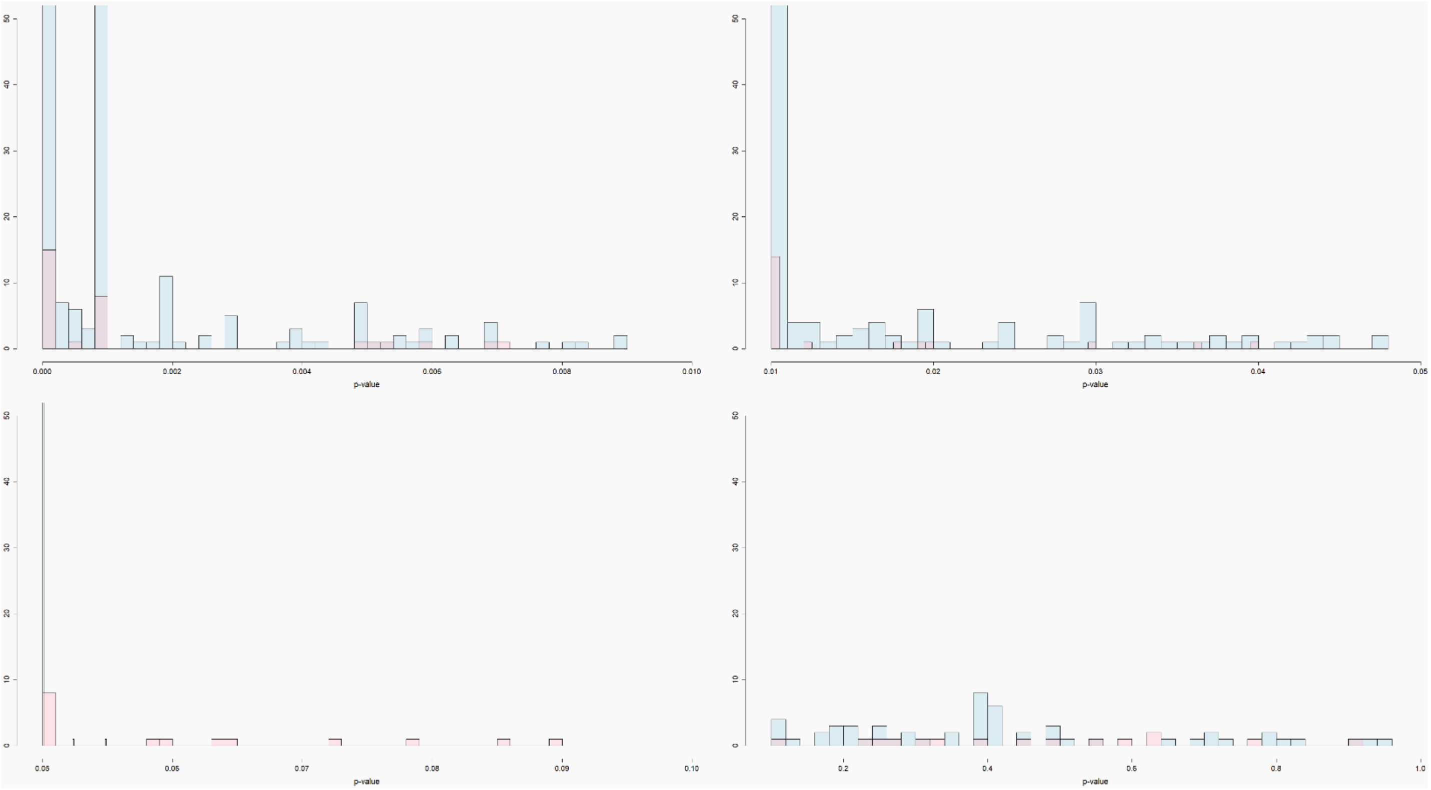

**Supplementary Figure 2.** Histogram of p-values from 720 experiments where p-values could be estimated. Blue represents p-values associated with increase in lifespan (positive SMDs) and pink represents p-values associated with decrease in lifespan (negative SMDs). Clockwise from upper left: 0<p<0.01; 0.01<p<0.05; 0.05<p<0.10; 0.10<p<1.0.

**Supplementary Table 1.** Comparison between mammal and non-mammal experimental results in the 36 compounds with both mammal and non-mammal experiments.

| Compound | # Mammal Experiments (Total N) | # Non-mammal Experiments (Total N) | Mammal SMD (p) | Non-mammal SMD (p) | Mammal % Increase in Lifespan | Non-mammal % Increase in Lifespan |
|---|---|---|---|---|---|---|
| Rapamycin | 12 (2100) | 6 (1919) | 0.33 (0.0052) | 1.84 (0.11) | 7.9% | 21.9% |
| Metformin | 5 (519) | 8 (3088) | 0.06 (0.80) | 0.42 (<0.0001) | 3.0% | 18.5% |
| Curcumin | 3 (293) | 8 (3869) | 0.45 (0.001) | 0.42 (0.004) | 7.2% | 8.4% |
| Vitamin E | 1 (47) | 10 (1621) | 0.57 (0.055) | 2.2 (0.17) | 12.5% | 6.5% |
| N-acetylcysteine | 1 (32) | 8 (4006) | 1.02 (0.0068) | 0.55 (0.0003) | 12.3% | 15.3% |
| Epigallocatechin gallate | 1 (60) | 7 (1692) | 0.52 (0.047) | 0.35 (<0.0001) | 11.2% | 16.9% |
| Quercetin | 1 (100) | 7 (3509) | -0.67 (0.0011) | 0.28 (<0.0001) | -9.4% | 9.5% |
| Aspirin | 3 (471) | 4 (880) | 0.07 (0.50) | 0.65 (0.017) | 1.0% | 16.2% |

| | | | | | | |
|---|---|---|---|---|---|---|
| Spermidine | 4 (360) | 3 (600) | 0.56 (0.0004) | 0.38 (<0.0001) | 9.0% | 22.0% |
| Melatonin | 3 (160) | 3 (344) | 1.07 (0.0041) | 0.83 (0.013) | 11% | 16.3% |
| Nicotinamide | 3 (370) | 3 (1365) | 0.25 (0.083) | 1.99 (0.24) | 3.1% | 19.4% |
| Alpha-keto glutarate | 1 (44) | 4 (769) | 0.83 (0.0088) | 0.49 (<0.0001) | 6.8% | 20.0% |
| Coenzyme Q10 | 3 (356) | 2 (394) | -0.27 (0.010) | 0.44 (<0.0001) | -6.5% | 17.7% |
| Nordihydroguaiaretic | 2 (455) | 3 (480) | 0.30 (0.37) | 0.58 (<0.0001) | 5.8% | 25.8% |
| Royal jelly | 1 (22) | 4 (2308) | 0.84 (0.060) | 0.48 (0.0041) | 17.9% | 14.1% |
| Alpha lipoic acid | 1 (177) | 3 (720) | 0.02 (0.91) | 0.42 (<0.0001) | 0.5% | 16.3% |
| Ascorbic | 1 (24) | 3 (404) | 0.94 (0.038) | 0.03 (0.91) | 19.6% | 2.0% |
| Epithalamin | 2 (171) | 2 (812) | 0.85 (0.0047) | 0.23 (0.0009) | 14.9% | 5.7% |
| Methionine | 2 (46) | 1 (300) | -0.62 (0.36) | -0.52 (<0.0001) | -8.3% | -6.3% |
| P-fluorophenylalanine | 2 (142) | 1 (120) | -0.31 (0.13) | 0.49 (0.0082) | -6.8% | 13.7% |
| Sodium borate | 1 (29) | 2 (400) | 0.15 (0.70) | -0.16 (0.60) | 0.9% | -6.0% |
| Aluminum chloride | 1 (28) | 1 (200) | -1.36 (0.0012) | -0.29 (0.043) | -7.7% | -4.7% |
| Berberine | 1 (44) | 1 (360) | 0.91 (0.004) | 0.06 (0.58) | 11.4% | 4.6% |
| Butylated hydroxytoluene | 1 (195) | 1 (500) | -0.51 (0.0005) | 0.31 (0.0005) | -11.8% | 13.0% |
| Calcium pantothenate | 1 (74) | 1 (197) | 0.47 (0.047) | 0.56 (0.0001) | 12.6% | 5.3% |
| D-glucosamine | 1 (146) | 1 (848) | 0.51 (0.0024) | 1.59 (<0.0001) | 5.5% | 2.7% |
| Dinitrophenol | 1 (60) | 1 (40) | 0.54 (0.042) | 0.81 (0.014) | 6.2% | 7.9% |
| Epitalon | 1 (108) | 1 (180) | 0 (0.98) | 0.24 (0.11) | -0.1% | 3.3% |
| Estradiol | 1 (370) | 1 (124) | 0.33 (0.0021) | 0.7 (0.0002) | 11.3% | 17.8% |
| Icariin | 1 (101) | 1 (321) | 0.25 (0.21) | 0.43 (0.0001) | 6.8% | 28.9% |
| Magnesium l-thiazolidine carboxylate | 1 (75) | 1 (331) | 0.45 (0.053) | 0.69 (<0.0001) | 9.1% | 21.4% |
| Oxaloacetate | 1 (310) | 1 (144) | 0.11 (0.32) | 0.42 (0.013) | 4.0% | 20.0% |
| Phenformin | 1 (54) | 1 (594) | 0.6 (0.033) | 0.27 (0.0011) | 11.6% | 28.5% |
| Simvastatin | 1 (682) | 1 (400) | 0.3 (0.0001) | 0.69 (<0.0001) | 19.0% | 12.6% |
| Skulachev quinone (SKQ) | 1 (50) | 1 (850) | 0.86 (0.0038) | 0.27 (0.0001) | 15.1% | 8.0% |
| Taurine | 1 (122) | 1 (470) | 0.8 (<0.0001) | 0.6 (<0.0001) | 14.5% | 20.4% |
